# COVID-19 Variants Database: A repository for Human SARS-CoV-2 Polymorphism Data

**DOI:** 10.1101/2020.06.10.145292

**Authors:** Allah Rakha, Haroon Rasheed, Zunaira Batool, Javed Akram, Atif Adnan, Jiang Du

**Author notes:** Corresponding authors Atif Adnan, Jiang Du. These authors contributed equally to this work (Allah Rakha, Atif Adnan).

## Abstract

COVID-19 is a newly communicable disease with a catastrophe outbreak that affects all over the world. We retrieved about 8,781 nucleotide fragments and complete genomes of SARS-CoV-2 reported from sixty-four countries. The CoV-2 reference genome was obtained from the National Genomics Data Center (NGDC), GISAID, and NCBI Genbank. All the sequences were aligned against reference genomes using Clustal Omega and variants were called using in-house built Python script. We intend to establish a user-friendly online resource to visualize the variants in the viral genome along with the Primer Infopedia. After analyzing and filtering the data globally, it was made available to the public. The detail of data available to the public includes mutations from 5688 SARS-CoV-2 sequences curated from 91 regions. This database incorporated 39920 mutations over 3990 unique positions. According to the translational impact, these mutations include 11829 synonymous mutations including 681 synonymous frameshifts and 21701 nonsynonymous mutations including 10 nonsynonymous frameshifts. Development of SARS-CoV-2 mutation genome browsers is a fundamental step obliging towards the virus surveillance, viral detection, and development of vaccine and therapeutic drugs. The SARS-COV-2 mutation browser is available at http://covid-19.dnageography.com.

## INTRODUCTION

The novel seventh strain of Coronaviridae associated with sever human to human transmission tentatively named 2019-nCoV that was officially declared a public health emergency of international concern by the World Health Organization (WHO) on January 31, 2020 [1]. Later on, the Coronavirus Study Group of the International Committee on Taxonomy of Viruses announced the name SARS-CoV-2 “Severe Acute Respiratory Syndrome Corona Virus 2” on 11 February 2020. Moreover, on the same day, the World Health Organization declared it a pandemic disease caused by the virus was “COVID-19” [2, 3]. The number of people infected with Coronavirus 19 will soon touch the five million mark in 209 countries including Pakistan. The first two cases of COVID-19 have been reported in Karachi and Islamabad by the Ministry of Health [4]. Although, COVID-19 is most prevalent in Pakistan as compare to other South East Asian countries [5]. To date, 27^th^May, 2,940,406 active cases, 357,894 (6.16%) deceased and 2,514,950 (43.26%) recovered has been reported worldwide [6].

The SARS-CoV-2 is the 3^rd^ most highly virulent strain of coronavirus that causes severe respiratory disease followed by SARS-CoV (severe acute respiratory syndrome Coronavirus) and MERS-CoV (Middle East respiratory syndrome coronavirus) with highest fatality rate [7, 8] while other members of this family 229E, OC43, NL63, HKU1 primarily involve in mild self-limiting disease such as “common cold” in the human population [9].

The first SARS-CoV-2 genome was sequenced and submitted to the NCVBI GenBank (NC-045512) on 12 January 2020 that consists of 29,903bp with close resemblance of SARS-CoV. Coronavirus recognized as highly evolving viruses, consists of non-segmented, positive-sense single-stranded RNA, with a high frequency of mutation and genomic recombination, [10, 11]. The virus genome is arranged in the order of 5’UTR (untranslated region), ORF1ab (open reading frame), 3’UTR, and nonstructural ORFs. ORF1ab encodes replicase polyprotein pp1a and pp1b, proteolytic cleavage of these two viral polyproteins produced 16nsp (nonstructural proteins) - Club shaped structural glycoprotein spikes(S) that gives the virus crown-like appearance-Envelope protein(E) - Membrane protein(M) - Nucleocapsid protein (N) [12].

The study on the hotspot of differences in the genome of SARS-CoV-2 is very important for investigating the etiology, epidemiology, prevention, and treatment of coronavirus infection. We aim to develop the SARS-CoV-2 mutation Browser v-1.3 that retrieved the full-length viral genome sequence data from National Center for Biotechnology Information (NCBI), Global Initiative on Sharing All Influenza Data (GISAID) and NGDC databases from all over the world to find the mutation or polymorphism in SARS-CoV-2 viral genome. This mutation browser will help the scientists and researchers to investigate the variation in the whole genome of SARS-CoV-2. Based on these findings, biomedical companies are also trying to develop rapid molecular diagnostic kits, vaccines, and therapeutic drugs for SARS-CoV-2 patients.

## Results

A total of 12,743 sequences collected from 105 regions (77 countries) were analyzed and filtered to acquire 8471 high coverage sequences containing 61917 mutations within 5415 mutating positions and an information browser was made available/accessible to the public. According to the translational impact, these mutations include 17,912 Synonymous mutations including 936 Synonymous Frameshifts and 33,823 Non-Synonymous Mutations including 17 Non-Synonymous Frameshifts.

Numbers of isolates with a certain number of mutations are also displayed in the browser and are given in **Table 1**. Most of the isolates with a count of 1047 have 6 mutations and terminals of the given range have the least number of mutations. This data is evident that the majority of sequences have lesser than ten mutations.

**Table 1.** Counters for Number of Isolates with certain number of Mutations.

| Number of Mutations | Number of Isolates |
| --- | --- |
| 0 | 75 |
| 1 | 126 |
| 2 | 181 |
| 3 | 185 |
| 4 | 321 |
| 5 | 910 |
| 6 | 1419 |
| 7 | 1495 |
| 8 | 1350 |
| 9 | 971 |
| 10 | 557 |
| 11 | 381 |
| 12 | 185 |
| 13 | 102 |
| 14 | 60 |
| 15 | 40 |
| 16 | 44 |
| 17 | 26 |
| 18 | 9 |
| 19 | 13 |
| 20 | 21 |
| Total | 8471 |

Mutation Browser is also furnished with a panel for range selection to visualize the number of variations within that particular range. Users can also directly select gene regions to visualize the number of mutations of transcribing genomic positions. An exemplary illustration of this visualization is given in **Figure 1**.

**Figure 1.**
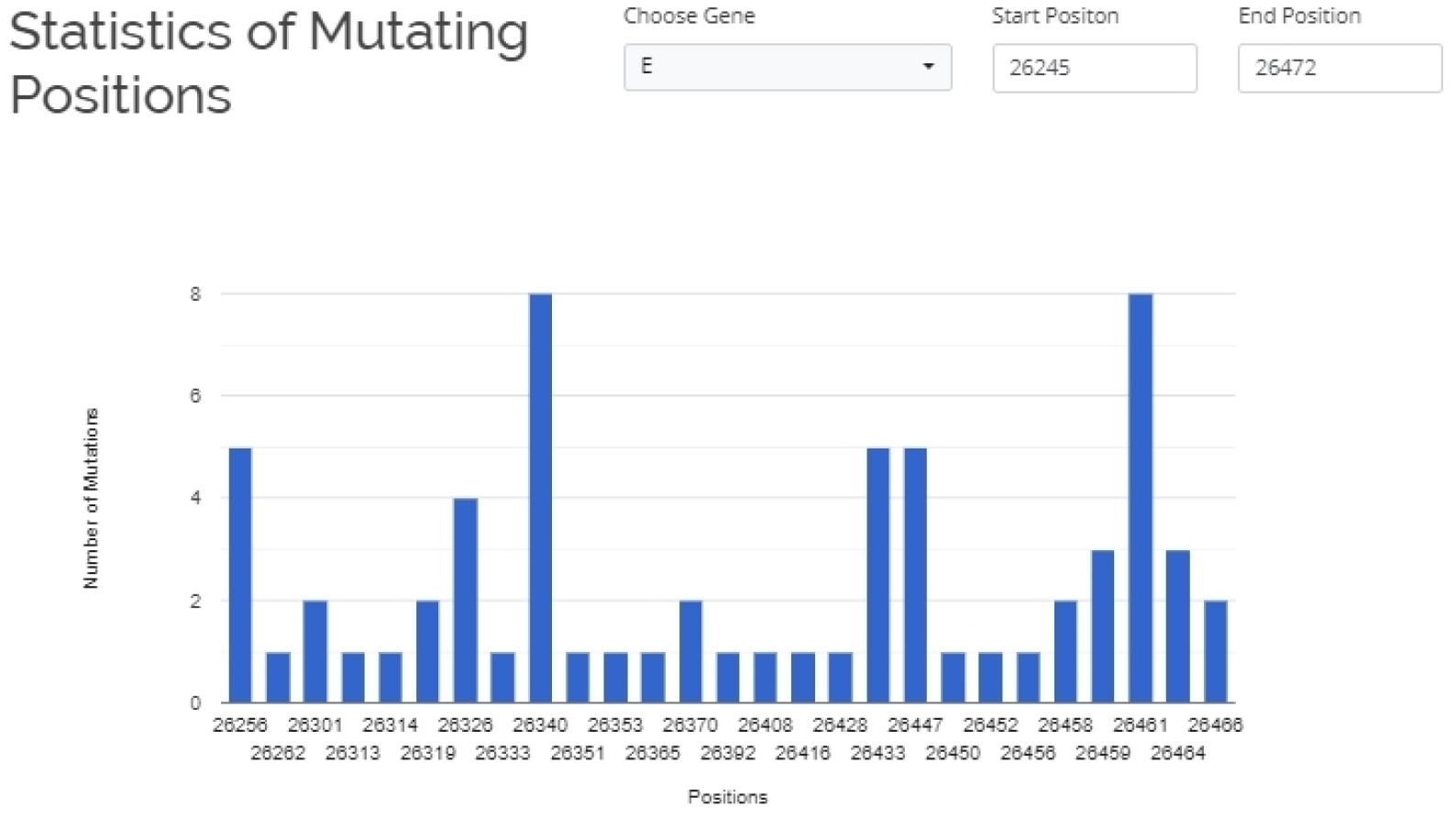
Statistics of Mutating Position Panel showing Number of Mutation over the Mutating Positions of E-Gene.

Another important tool in the mutation browser is “Primer Infopedia” where the incorporation of available primers for COVID-19 sequence amplifications to select their regions directly from the list and visualize the mutation details of those regions along with the details of geographical regions curated from sequence information. This utility helps a lot for a quick review of the reliability of the primers for the SARS-CoV-2 genome.

## Discussion

Healthcare professionals and evidence-based medical experts of different countries launched the COVID-19 information portal which aggregates immediate information updates related to COVID-19 pandemic from reliable sources (Ref). These COVID-19 dashboards and portals provide general information about disease epidemiology, symptoms, diagnosis, preventive measure, medication as well as morbidity and mortality ratio [13, 14]. Along with these general COVID-19 portals, there was also dire need to launch a diagnostic-based portal that facilitates the researchers and scientists to develop the state of art molecular detection assays, vaccines, and therapeutic drugs against COVID-19. In the current situation of the COIVD-19 pandemic, we have developed and launched the SARS-CoV-2 mutation Browser v-1.3 on date 28-Feb-2020 that is updated regularly.

The SARS-CoV-2 mutation browser is easy, efficient, and user friendly that only deals with novel coronavirus and display the viral genome variation around the globe on a single platform for the convenience of the researchers and virologists through World Wide Web http://covid-19.dnageography.com. The browser features comprehensive integration of a genomic sequence of SARS-CoV-2along with their metadata information from NGDC, NCBI, and GISAID. Furthermore, it is open access mutation browser that displays the region and gene-wise (5UTR, ORFab, S, E, M, N, 3UTR) distribution of mutation data of particular virus isolate and also provides position and statistics of mutations. Moreover, viral primer sequence information is also displayed on the same page “primer infopedia”. The mutation browser has close resemblance with a Chinese“2019nCOVR” **2019 Novel Coronavirus Resource** browser with additional functions like it deals with all coronavirus family, retrieved the viral genome sequence data from NCBI, GISAID, NMDC and CNCB/NGDC [15].

The evolution rate of coronavirus could be 10^−4^ substitute per bp per year as a typical RNA virus therefore; the mutation could be occurring during each replication cycle [9]. However, in the present bio-information browser, 5688 SARS-CoV-2 viral sequences from ninety-one regions were examined and a total of 39920 mutations were found. Importantly, the database showed position and type of mutation, consequence, and consequence type in the SARS-CoV-2 genome. Therefore, to avoid false-negative results, data from this database will be helpful to design the PCR primers and probes with maximum specificity and sensitivity for the diagnosis of COVID-19. A potential limitation of this mutation browser is that since all SARS-CoV-2 genome sequences used in this study were retrieved from different databases and the accuracy of these genome sequences could not be verified although these partial or complete sequences have been published worldwide.

In conclusion, the COVID-19 mutation browser is a worthful addition to the currently available coronavirus resources that will allow the researchers to access data expeditiously and intuitively on SARS-CoV-2 mutation. We expect the SARS-CoV-2 mutation browser will assist the process of study design and be a key resource for the COVID-19 community.

## Materials and Methods

### Study Design

#### Data Collection and Multiple Sequence Alignment

A total of 12,743 high coverage complete genomes of SARS CoV-2 isolates from 77 countries (105 regions) were retrieved from GISAID.org [16]. Complete detail about the origin of each sequence is given in ***Supplementary Table 1***. The reference genome sequence of the virus was also retrieved from NCBI-GenBank (GenBank: MN908947.3) [17, 18]. All of these viral genome sequences were aligned with reference sequence in different sets by using Clustal Omega Server provided by EMBL-EBI [19, 20].

#### Processing of Aligned Sequences

The aligned sequences were processed by an in-house python script to extract mutations and filter sequences. Based on a filter over the number of differences, only high-quality sequences with a max of 20 differences were selected for the database. Our python script could only call all the variants except deletions at terminal regions of aligned sequences and where nucleotide changes to “N” (Unknown nucleotide). This script was also programmed to select and curate mutations only from those sequences having the utmost 20 mutations and no addition greater than two base pairs.

#### Annotation of Nucleotide Mutations

All the nucleotide variations were also annotated for their translational consequences along with consequence types. For achieving this milestone, an in-house built database was generated having the annotations of each known consequence permutation and then a python script was written to annotate each mutation and curate its consequence and consequence type.

#### Development of Mutation Browser

The SARS-CoV-2 Mutation Browser v.1.3 has been generated the viral genome sequences data with different types of mutations. Two viral genome sequences of Pakistani isolates retrieved from NCBI were also processed through the same procedure and are provided separately on the browser. This browser is coded using modern programming languages including HTML5, JavaScript, JQuery, PHP, and equipped with MySQL server for rapid retrieval of information. This browser is designed to provide the user with visualizations of region-wise differences, Gene wise differences, Hotspot of differences in Genome, Consensus sequence, and all differences of a sample from reference genomes and their consequence along with summarized information represented in tabulated format. This portal is also coded to show some *basic* stats of isolates regarding several mutations and genomic positions concerning the number of mutations with the integration of Google Charts.

#### Primer Infopeida

All the primers reported for SARS-CoV-2 diagnostics and research purposes were manually curated from literature and websites. All these primers were aligned to the reference sequence using the NCBI BLAST tool to select fully reference mapping primers and curate them with their regions according to the reference sequence. Mutations Database and Primers Database were then integrated into another “Primer Infopedia” tool that is also a part of SARS-CoV-2 Mutation Browser v.1.3.

## ACKNOWLEDGEMENTS

We acknowledge University of Health Science, Lahore and China Medical University for providing computational resources and financial support for this project.

## ETHICS DECLARATIONS

The study was approved by the ethical review board of University of Health Sciences (UHS) Lahore, Pakistan in accordance with the standards of the Declaration of Helsinki.

## AUTHOR CONTRIBUTIONS

A.R. J.A and A.A developed the idea. A.R and A.A. wrote the manuscript, A.R., H.R., A.A., D.J., and J.A. performed dry lab work. A.R. and A.A. modified the manuscript. All authors reviewed the manuscript.

## COMPETING INTEREST

No potential conflict of competing interest was declared by the authors.

## Notes

### Competing Interest Statement

The authors have declared no competing interest.

